# A sulfonated cartilage interpenetrating polymer network reinforces and protects the extracellular matrix of degraded cartilage

**DOI:** 10.1101/2025.07.28.667232

**Authors:** Christian D. DeMoya, Pierson Husted, Dev R. Mehrotra, Michael B. Albro, Brian D. Snyder, Mark W. Grinstaff

**Author notes:** Corresponding authors: Mark W. Grinstaff, Brian D. Snyder, Michael B. Albro.

## Abstract

Cartilage extracellular matrix (ECM) comprises a type-II collagen fibril network that affords structure and tensile strength, complemented by a negatively charged, sulfated glycosaminoglycan (GAG) matrix that retains interstitial water. These components act synergistically, bestowing the rheological and tribological material properties essential to cartilage function. At the onset of osteoarthritis, a disease characterized by cartilage degeneration, GAGs diminish from the ECM reducing interstitial fluid load support (*IFLS*) and transferring load to the collagen fibril network, which subsequently breaks down, culminating in increased hydraulic permeability, and decreased cartilage stiffness. We restore the material properties of damaged cartilage critical to diarthrodial joint function by forming an interpenetrating polymer network (IPN) with the native collagen using a synthetic, hydrophilic, and biocompatible GAG-mimetic polymer. Upon visible light activation, the monomer, 3-sulfopropylmethacrylate (SPM), and the crosslinker, polyethylene glycol diacrylate (PEGDA), form a sulfonated and anionic IPN that entangles and fills the existing porous degraded collagen matrix. Mechanistically, the highly sulfated, anionic SPM IPN retards water transport, reestablishes collagen fibril network integrity, and restores tissue *IFLS*, thereby returning the stiffness and viscoelastic properties of degraded cartilage to healthy levels. Additionally, the SPM IPN protects cartilage from further degradation by reducing the infiltration of inflammatory cytokines that upregulate catabolic matrix metalloproteinases and downregulate GAG production.

**Statement of significance:** Amelioration of OA requires a comprehensive approach: neutralize or impede catabolic enzymes that degrade cartilage and reconstitute damaged cartilage by augmenting tissue ECM constituents. Currently, there are no clinical treatments that restore the viscoelastic material properties of hyaline cartilage tissue critical to its mechanical function and impart chondroprotection after OA induction. This work suggests that reconstituting GAG-depleted cartilage using a synthetic sulfonated interpenetrating polymer to reestablish *IFLS* that can be instilled into the joint and polymerized with white light during conventional arthroscopy represents a novel, minimally invasive, clinical treatment for early OA.

## 2. Introduction

Hyaline cartilage is the smooth, hydrated viscoelastic, biphasic, composite material covering articular joint surfaces. The extracellular matrix (ECM) comprises a type-II collagen (COLII) fibril network (5-20 w/v%) that provides tensile strength, complemented by a porous-permeable glycosaminoglycan (GAG) matrix (5-15 w/v%) of negatively charged, sulfated polysaccharides that regulate the transport of interstitial water (70-90 w/v%).[1] These constituents act synergistically imparting the rheological and tribological material properties essential for articular cartilage function: support applied compressive and shear loads, while maintaining a nearly frictionless interface between articulating joint surfaces. Essential to cartilage function is the ability of the ECM to coordinate water via highly sulfated, anionic, GAGs decorating a core protein that aggregates to hyaluronic acid, forming a bottlebrush structure. Pressurization of entrapped interstitial fluid provides interstitial fluid load support (IFLS) [2, 3] critical to sustaining applied compressive joint loads. Compression deforms the COLII network, decreasing pore volumes while relatively increasing the local fixed negative charge density that act together to retard interstitial fluid transport. This strain-dependent decrease in tissue permeability increases the viscous drag of fluid flowing through the porous COLII network responsible for cartilage’s non-linear, viscoelastic behavior.[4] Besides affording load support, the interstitial fluid also contributes to elastohydrodynamic joint lubrication.[3-9] Compressive deformation induces fluid efflux from the hyaline cartilage, creating an interposed fluid film between the articulating joint surfaces which decreases the coefficient of friction (COF) and protects the tissue from mechanical shear wear. The COF inversely correlates with the *IFLS*, achieving its lowest value when *IFLS* is greatest.[8, 10]□ Aside from contributing to cartilage’s rheological and tribological material properties, the semipermeable GAG-rich ECM is chondroprotective, restricting infiltration of damaging macromolecules (e.g., anionic cytokines and matrix metalloproteases) into the tissue from the synovial fluid by the excluded volume effect [11, 12].

Osteoarthritis (OA) in its many forms represents an incapacitating, chronic, non-inflammatory, synovial joint arthrosis that vexes >520 million adults world-wide. It manifests clinically as pain, swelling and decreased joint mobility with associated treatment costs approaching $486.4 billion annually.[13] Various biochemical,[14-17] biomechanical [18-20], metabolic [21-25], and gene-regulated [26-28] mechanisms, occurring over an extended timeline, contribute to OA pathophysiology. Injurious mechanical overloading of joint tissues because of trauma, internal derangement, joint instability, ligamentous deficiency, skeletal malalignment, and/or obesity provokes an inflammatory cascade that affects all synovial joint tissues and, in particular, hyaline cartilage [29, 30]. Early in the disease process, GAGs are depleted from the cartilage because of cytokine-mediated upregulation of matrix metalloproteinases and downregulation of GAG production [31-33]. Depletion of cartilage GAG reduces *IFLS*, transferring load to the COLII fibril network, which subsequently breaks down, culminating in increased hydraulic permeability, and decreased cartilage stiffness [34]. Increased tissue porosity and decreased GAG content reduces the excluded volume effect through charge interactions, diminishing repulsion of anionic solutes [35] allowing proinflammatory cytokines in the synovial fluid to permeate the GAG-depleted ECM and access chondrocytes.

Amelioration of OA requires a comprehensive approach: 1) correct the mechanical factors contributing to acute or chronic chondrocyte injury and tissue wear; 2) abrogate the inflammatory cascade and neutralize catabolic enzymes that degrade cartilage; and, 3) reconstitute damaged cartilage material properties by augmenting tissue ECM constituents. *Currently, no clinically used pharmacological treatments mitigate cartilage wear and impart chondroprotection after the induction of OA*. Conventional tissue engineering strategies rely on excising the damaged cartilage and replacing it with an engineered neocartilage implanted into the defect using cell-free or cell-laden hydrogel constructs comprised of natural or synthetic polymers [36-40] composed of poly (ethylene glycol),[41, 42] poly (vinyl alcohol),[43] poly (lactic acid),[44, 45] hyaluronic acid,[46, 47] alginate,[48, 49] or agarose[50, 51]. An alternative approach salvages the existing cartilage ECM by restoring the material properties of the damaged cartilage critical to diarthrodial joint function by forming an interpenetrating polymer network (IPN) with native COLII using a synthetic, biocompatible, anionic, hydrophilic, GAG-mimetic polymer to reestablish *IFLS* and reestablish the integrity of the collagen fibril network. IPNs are known to enhance the material properties of engineered hydrogels [52-55]. Our group has developed synthetic zwitterionic polymers that integrate with the existing collagen network [56-58]: a reactive monomer, crosslinker, and photo-initiator are introduced into the GAG depleted ECM; light activated polymerization installs the polymer network throughout the native fibrillar COLII network which resurrects the biomechanical and transport restrictive properties of damaged cartilage. We present a new anionic IPN that mimics the naturally occurring proteoglycan network accomplished by combining 3-sulfopropylmethacrylate potassium salt (SPM), poly (ethylene glycol-diacrylate) (PEGDa) and a photoinitiation system consisting of triethanolamine (TEOA), n-vinylpyrrolidone (NVP), and eosin Y. Analogous to naturally occurring GAGs, the SPM imparts anionic charge via a sulfonate group, while the crosslinks provided by PEGDa ensure that the IPN interlinks with the existing collagen matrix (**Fig. 1**). The photoinitiation system provides temporal control over the polymerization process, ensuring sufficient time for the components of the IPN to diffuse into the damaged cartilage ECM and has been used safely *in vivo* and in the clinic [59-61]. The o*bjective* of this work is to identify the mechanism by which the SPM IPN augments GAG depleted cartilage ECM. We hypothesize that SPM IPN reconstitutes the material properties of degraded cartilage by filling the GAG depleted porous ECM with a dense hydrogel that coordinates water and restores IFLS thereby improving tissue stiffness, prolonging tissue relaxation, and slowing solute permeability critical to hyaline cartilage function.

**Figure 1.**
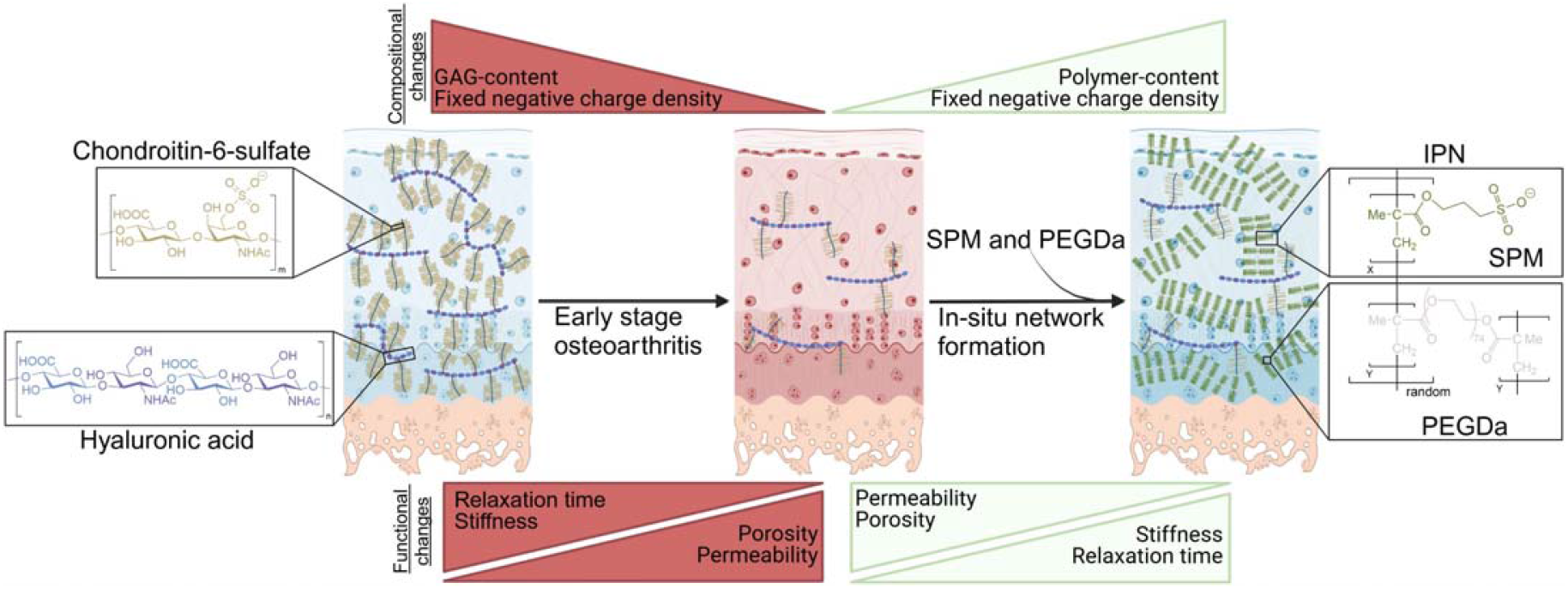
Healthy cartilage contains an extensive proteoglycan network, consisting of a core protein decorated with GAGs, mainly chondroitin sulfate, that is then tethered to high molecular weight hyaluronic acid chains. Depletion of the GAG content in OA leads to impaired cartilage function and perpetuates a cycle of degradation. *In situ* formation of a cartilage IPN using SPM and PEGDa replicates key characteristics of the natural GAG network to restore the properties of the cartilage to healthy levels.

## 3. Materials and methods

### 2.1 Reagents

PEGDa (Average M_W_ 3,400 Da), and Antibiotic-Antimycotic (100X, anti-anti) were acquired from Fisher Sci (Hampton, NH). Recombinant human interleukin-1α (IL-1α), and IL-6 were acquired from R&D Systems Inc (Minneapolis, MN). ELIZA kits against IL-1α (ab178008), and IL-6 (ab178013) were acquired from abcam (Cambridge, UK). SPM, TEOA, NVP, eosin Y, papain, fluorescein-isothiocyanate conjugated dextran (FITC-dextran, M_W_ 70 kDa), chondroitinase-ABC, D-(+)-maltose monohydrate, 1,9-Dimethyl-Methylene Blue (DMMB), benzamidine hydrochloride (BHCl), and ethylenediaminetetraacetic acid (EDTA) were acquired from MilliporeSigma (Burlington, MA). All reagents were used as provided by the vendor.

### 2.2 Immature bovine and mature bovine cartilage harvest

Healthy cartilage plugs (Ø=2 mm, thickness=1 mm) were harvested from the middle and deep zone of the femoral groove of 2-month-old calves (Green Village Packing Co, NJ) and stored frozen at -20º C in 400 mOsm saline containing protease inhibitor BHCl (5 mM), anti-anti (1X), and calcium ion chelating agent EDTA (5 mM). Osteochondral plugs (∼5 mm X 4 mm) were harvested from a visibly degenerated medial tibial plateau of a skeletally mature bovine stifle joint (Animal Technologies, Inc, Tyler, Texas) along a rectangular grid using a fine-tooth handsaw. The subchondral bone was cut parallel to the articular surface to create a plano-parallel surface for mechanical testing.

### 2.3 Experimental plan

We began by assessing the biochemical composition of the healthy, GAG-depleted, and IPN-treated cartilage samples; specifically, the GAG content, water content, and IPN content. To assess the ability of the IPN to restore the mechanical properties of GAG-depleted cartilage to those of healthy cartilage, a repeated measurement study was conducted with mechanical testing performed on cartilage samples in healthy, GAG-depleted, and IPN-treated conditions. Additionally, to identify the IPNs ability to induce Donnan osmotic swelling pressure in a similar manner as the native GAG network, healthy and IPN-treated cartilage plugs underwent a repeated measure study with mechanical testing done in solutions of variable ionic strength. Finally, we determined the ability of the IPN to restrict solute infiltration into the ECM of cartilage samples through an independent measure study conducted on healthy, GAG-depleted, and IPN-treated cartilage plugs.

### 2.4 GAG depletion of immature cartilage

To create a consistent model of GAG-depleted cartilage, healthy immature bovine cartilage was treated with chondroitinase-ABC. After thawing cartilage plugs to room temperature, the plugs were incubated in chondroitinase-ABC (0.1 U/mL in 50 mM tris(hydroxymethyl)aminomethane, 60 mM sodium acetate, 0.02% bovine serum albumin, pH 8.0) at 37º C for 24-hours to deplete healthy cartilage of GAG. Residual enzyme and degraded GAGs were removed by washing the plugs with 400 mOsm saline.[62]

### 2.5 Water content and GAG quantification

To calculate water content, the wet and dry weights of all plugs were determined from gravitational measurements before and after lyophilization. GAG content was quantified by digesting plugs with papain (1 mg/mL) at 60º C for 24 hours and then determining the GAG concentration in the lysates by DMMB colorimetric assay.

### 2.6 IPN instillation into cartilage

The IPN solutions were made by dissolving SPM at the appropriate concentration (5 w/v%, 20 w/v%, 40 w/v%, and 60 w/v%) in 400 mOsm saline, along with PEGDa (0.01 mol PEGDa/ mol SPM), TEOA (115 mM), NVP (94 mM), and eosin Y (0.1 mM). Samples were incubated in IPN solutions protected from the light for 24-hours at room temperature on an orbital shaker. Plugs were removed from the IPN solution, blotted dry, and photo-irradiated in a humidity chamber with high intensity white light using a HL150-A halogen illuminator (AmScope, Irvine, CA) for 15-minutes and washed with saline to remove unreacted material. The cartilage samples were then stored in 400 mOsm saline.

### 2.7 Quantification of IPN content

After measuring the water content of IPN-treated cartilage, IPN content was quantified by digesting plugs with papain (1 mg/mL) at 60º C for 24 hours; the lysates were dialyzed against deionized water using a 50 kDa molecular weight cutoff dialysis membrane (Repligen, Waltham, MA) for 48-hours to remove proteinaceous contaminants in solution. The remaining lysate was lyophilized, and the weight of the residual polymer determined by gravitational measurement.

### 2.8 Mechanical testing of cartilage

Material properties were derived using a customized uniaxial servo-driven actuator equipped with a stainless-steel platen and in-line 500 g load cell. The diameter and thickness of each plug was measured using a digital caliper. Chondral plugs were submerged in test solutions (400 mOsm saline, 0 mOsm saline, 2000 mOsm saline, or 400 mM maltose solution) for the duration of the mechanical procedure and were subjected to an initial 50 g creep load for 300 s followed by a 4-step incremental stress relaxation protocol where a 5% incremental strain was applied at a rate of 8 µm/s over each step. Using established procedures [63] a MATLAB script was used to calculate the results. The peak stress (σ_peak_) and equilibrium stress (σ_eq_) at each step were identified. The equilibrium modulus (E_eq_), which characterizes the non-fibrillar component of the cartilage stiffness, was obtained from the slope of the best fit line to σ_eq_versus strain (**Fig. S1**). The instantaneous modulus (E_inst_) at each step was calculated from σ_peak_divided by the corresponding strain. The initial E _inst_ 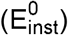 was determined from the y-intercept and strain-dependent E _inst_ 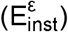 derived from the slope of the best fit line of E _inst_ versus strain (**Fig. S1**).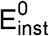 describes the modulus of the fibrillar component contributing to tissue stiffness at the onset of the strain application, while 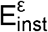 describes the magnitude of the change in the modulus of the fibrillar component contributing to tissue stiffness as the strain changes.The relaxation curve at each step was fit to a stretched exponential model, 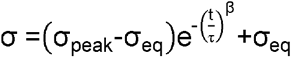 allowing computation of the relaxation time (τ) and stretching parameter (β).[64]

### 2.9 Test of healthy and IPN-treated cartilage in solutions of variable ionic strength

To demonstrate that SPM augments the fixed negative charge density of cartilage ECM, the Donnan effect of bathing solution ion concentration and osmolarity on cartilage material properties was evaluated on intact and SPM IPN augmented GAG-depleted cartilage plugs. Mechanical testing in 400 mOsm saline reflects both the Donnan effect of Na^+^ cations attracted to the fixed negative charge components (native GAGs and SPM) in the cartilage ECM and the effect of bathing solution osmolarity. Plugs mechanically tested in a water bath (0 mOsm) after 24-hours of equilibration in water reveal how the non-permeable anions (GAG and SPM) effect tissue osmotic pressure as ions equilibrate within and outside the tissue. Plugs equilibrated in charge neutral 400 mM maltose osmotic solution (osmolarity equivalent to healthy synovial fluid) for 24-hours before mechanical testing portray the effect of osmolarity alone on tissue material properties, devoid of diffusible cations. Additionally, the non-electrostatic contributions of the GAG network and IPN to the mechanical properties of healthy and IPN-treated cartilage, respectively, were determined by conducting mechanical testing of the samples in high ionic strength solutions. Specifically, following testing in 400 mOsm saline, the cartilage samples were equilibrated in 2000 mOsm saline solutions for a minimum of 24-hours with 3 solution changes. The cartilage then underwent the previously described 4-step stress relaxation procedure to measure the mechanical testing.

### 2.10 Solute infiltration into cartilage samples

The slowing of solute permeability into cartilage after tissue augmentation with SPM IPN was assessed by measuring the infiltration of different solutes into the plugs. FITC-dextran, IL-1α, and IL-6 were used as the solutes of interest and their physical properties are summarized in table S1. Plugs were submerged in an absorption bath of known solute concentration (FITC-dextran 1 mg/mL, IL-1α 50 ng/mL, and IL-6 0.2 ng/mL) for 24-hours at room temperature with shaking, then transferred to a saline desorption bath for an additional 24-hours at room temperature with shaking. Specimens were protected from light during both incubation periods. Solute concentrations in the absorption and desorption baths were measured using a spectraMax iD3 plate reader (Molecular Devices, San Jose, CA). The concentration of FITC-dextran was determined against a FITC-dextran standard curve. IL-1α and IL-6 concentrations were measured using ELIZA kits according to manufacturer’s protocol. Partition coefficients (K) were calculated: K= V_D_×C_D_/V_C_×(C_A_-C_D_), where: V_D_ is the volume of the desorption bath, C_D_ is the concentration of the desorption bath, C_A_ is the concentration of the absorption bath, and V_C_ is the volume of the cartilage plug. Following incubation in the adsorption bath, fluorescent images of FIT-dextran infiltrating into cartilage plugs were obtained. The plugs were blotted dry, cut in half, mounted on a glass slide and imaged using an FV3000 laser scanning confocal microscope (Olympus, Tokyo, JP) to visualize tissue penetration.

### 2.11 Indentation testing

To measure stress relaxation, osteochondral plugs submerged in a reservoir of 400 mOsm saline were subjected to incremental deformation applied by a servo-driven uniaxial actuator under displacement control via a 3 mm spherical indenter. The tissue was indented to a depth of 100 µm at a rate of 2.5 µm/s and allowed to relax for 180 s. Assuming normal Hertzian contact, the modulus (E) was determined by fitting the force-displacement curve to the Hertz model of ball-on-a-flat-surface for indentation using the equation 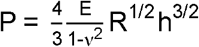, where P is the applied load, ν is Poisson’s ratio, R is the tip radius, and h is the indentation depth.[65]

### 2.9 Raman compositional assessment of mature bovine cartilage

Raman spectroscopy provided a non-destructive, optically based, label free method to monitor the composition of the cartilage ECM. Raman spectra were acquired using a custom Raman probe comprised of a hollow needle tip with a distally mounted Ø2 mm diameter sapphire ball lens, coupled to an NIR diode laser (ex=785 nm; 100 mW output; IPS, Plainsboro, NJ) and an Eagle Raman-S spectrometer (Ibsen Photonics, Farum, DK) equipped with a NIR optimized charge coupled device (Ivac, Andor, Belfast, Ireland). For Raman spectral acquisitions, the probe tip was positioned perpendicular to the articular surface and spectra were captured over a 30 s integration time. Following subtraction of the probe background noise, a 3^rd^ order polynomial baseline correction and subsequent area-under-curve normalization was applied to the 800-1800 cm^-1^ spectral “fingerprint” region. The resulting cartilage “fingerprint” spectral profile, Cartilage_spectra_was decomposed according to the multivariate linear regression model: [66, 67]

Cartilage_spectra_=GAG_score_×GAG_REF_+Collagen_score_×Collagen_REF_+Water_score_×Water_REF_+Bone_score_×Bone_REF_+SPM_score_×SPM_REF_ where the derived regression coefficients GAG_score,_ Collagen_score_, Water_score_, Bone_Score_ and SPM_score_ reflect the relative contribution of each individual constituent to the composite spectra; GAG_REF_, Collagen_REF_, Water_REF_, Bone_REF_ and SPM_REF_ are reference spectra measured from purified chemicals of each constituent: GAG_REF_= chondroitin sulfate (bovine cartilage; C6737 Sigma); Collagen_REF_= type-II collagen (chicken sternal cartilage; C9301 Sigma); Water_REF_= PBS; Bone_REF_= Mature bovine subchondral bone; SPM_REF_ = purified polymer synthesized outside of a cartilage plug. Additionally, the high wavenumber spectral range (2800-3800cm^-1^) was preprocessed using a linear baseline subtraction and area-under-curve normalization. The resulting spectral profile was analyzed for the area under the organic-content-associated CH_2_ spectral region (CH_area_) and area under the water-content-associated OH region (OH_area_).[68]

### 2.10 Statistical analysis

Statistical and graphical analyses were conducted using GraphPad Prism 10.0 software (GraphPad Software, Inc., San Diego, CA, USA). All data are represented as means and standard deviations (mean±S.D.). Experiments in which the same sample underwent repeated testing in different conditions used paired statistical tests. Significance was determined with an α-value of 0.05.

## Results

### *3*.*1 In situ* photopolymerization of 3-sulfopropylmethacrylate forms a cartilage IPN

Chondroitinase-ABC efficiently depletes healthy immature bovine cartilage of native GAG from 5.17±0.91% to 0.25±0.46% (**Fig. 2A**). The IPN solution effectively polymerizes with photoirradiation using high intensity white light as confirmed by ^1^H NMR (**Fig. S2**). SPM partions into the GAG-depleted cartilage matrix across increasing SPM monomer bath concentrations of 5 w/v%, 20 w/v%, 40 w/v% and 60 w/v%, followed by photoirradiation to interlink the polymer with the existing collagen matrix, achieving tissue IPN contents of 5.1±2.58 w/v%, 6.6±2.83 w/v%, 7.9±4.27 w/v% and 9.1±3.86 w/v%, respectively **(Fig. 2B)**

**Figure 2.**
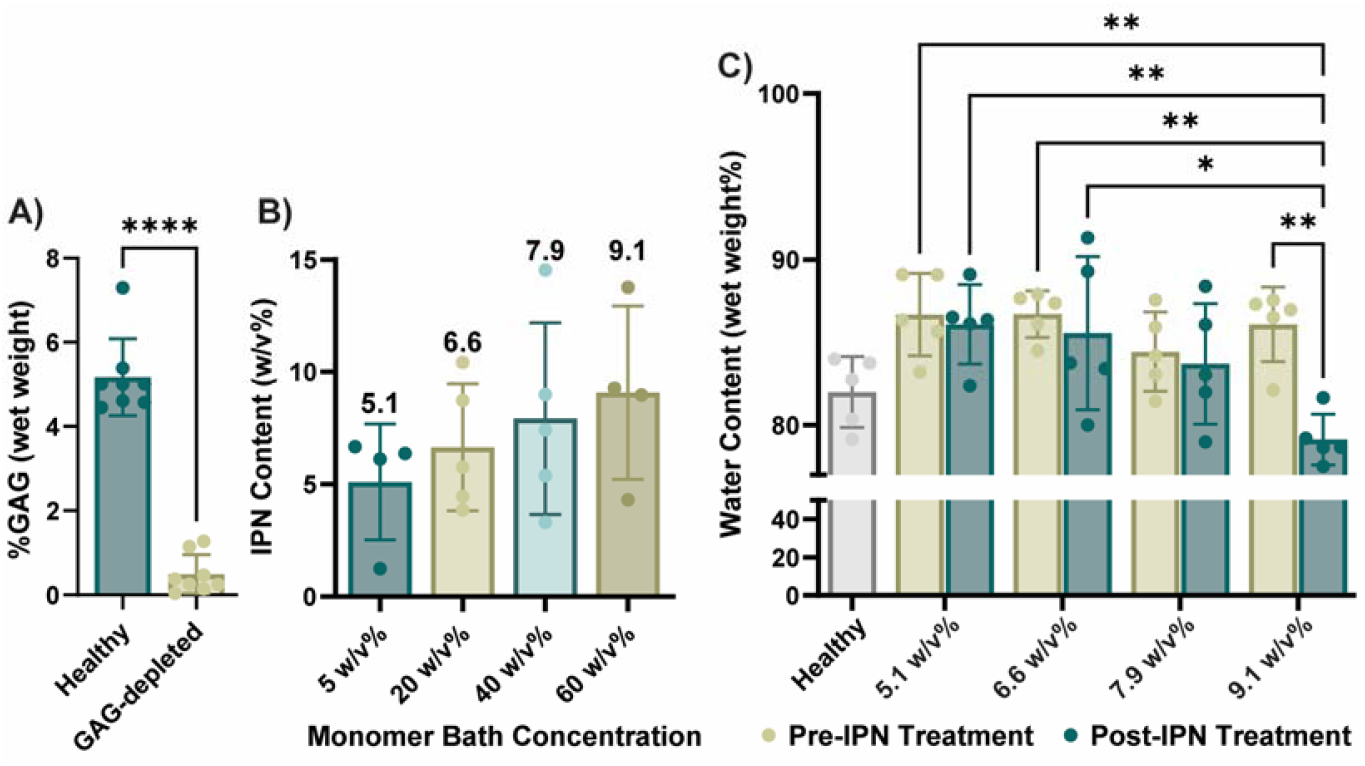
Characterization of the GAG, IPN, and water content across treatment groups. **A)** The GAG-content of chondroitinase-ABC treated cartilage is significantly less than intact cartilage. n=8 cartilage plugs per group. Data is represented as mean±S.D. Significance determined by unpaired t-test, ^****^ p<0.0001. **B)** GAG-depleted cartilage was incubated in four different monomer bath concentrations for 24-hours followed by photoirradiation to form the IPN. The resulting amount of polymer present in the cartilage was calculated and found to increase with increasing monomer bath concentrations. n=4 or 5 cartilage plugs per group **C)** The water content of intact cartilage and GAG-depleted cartilage, before and after IPN treatment was measured in n=5 cartilage plugs per group. The water content of cartilage increases with GAG depletion and is returned to that of intact cartilage only with 9.1 w/v% IPN. Data is represented as mean±S.D. Significance determined by 1-way unpaired analysis of variance (ANOVA), ^*^ p<0.05, ^**^

Interstitial water content reflects cartilage tissue porosity, therefore we measured the water content in healthy, GAG-depleted, and IPN-treated cartilage at these four concentrations of IPN treatment. Compared to healthy cartilage water content (82.0±2.15%), after GAG-depletion, and before IPN formation, the water content of the corresponding cartilage plugs increases to 86.69±2.50%, 86.72±1.42%, 84.44±2.39%, and 86.09±2.26% for the 5.1 w/v%, 6.6 w/v%, 7.9 w/v%, and 9.1 w/v% IPN groups, respectivley (**Fig 2C**), indicating that tissue porosity increases after enzymatic GAG depletion for all groups prior to IPN induction. After incubation in the SPM bath and photopolymerization to form cartilage IPN concentrations of 5.1 w/v%, 6.6 w/v% and 7.9 w/v%, the water content of the corresponding IPN treated cartilage plugs remains elevated compared to healthy cartilage: 86.10±2.41%, 85.56±4.64%, 83.70±3.65%, respectively. Only the 9.1 w/v% IPN group achieves an intact cartilage water content of 79.14±1.54% **(Fig. 2C**). This suggests that an SPM bath concentration of 60 w/v% is required to form a sufficient IPN in the porous-permeable GAG-depleted ECM and that the infiltrated SPM-IPN reduces tissue porosity by insinuating into the pores.

### 3.2 Restoration of Material Properties Depends on IPN Concentration in GAG-depleted Cartilage

We measured the elastic and viscoelastic material properties of healthy, GAG-depleted, and IPN-treated cartilage using a 4-step stress relaxation procedure. For the four groups, prior to IPN infiltration, GAG depletion decreases E_eq_(0.075±0.064 MPa-0.099±0.072 MPa) by 75% compared to that of the healthy cartilage (0.30±0.12 MPa - 0.38±0.11 MPa) (**Fig. 3A**). The E_eq_marginally increases after photopolymerization at the three lowest IPN concentrations; IPN tissue concentrations of 5.1 w/v%, 6.6 w/v% and 7.9 w/v% achieved E_eq_of 0.10±0.08 MPa, 0.11±0.11 MPa, and 0.12±0.1 MPa, respectively. However, 9.1 w/v% IPN augments the more than 8-fold to 0.84±0.44 MPa, exceeding that of the healthy cartilage (**Fig. 3A**). For the four groups prior to IPN infiltration, GAG-depletion reduces cartilage by ∼60% (1.35±0.93 MPa - 2.24±0.83 MPa) relative to that of healthy cartilage (4.05±1.86 MPa - 5.21±1.90 MPa, **Fig. 3B**). restores to that of the intact cartilage only for the 9.1 w/v% IPN group (3.85±1.26 MPa). Similarly, for the four groups prior to IPN infiltration, GAG- depletion reduces by 42% (15.83±5.26 MPa – 19.41±5.74 MPa) compared to that of the healthy cartilage (27.06±7.41 MPa – 34.05±10.02 MPa, **Fig. 3C**). After photopolymerization, IPN tissue concentrations of 5.1 w/v%, 6.6 w/v% and 7.9 w/v% do not restore (15.71±4.26 MPa, 18.48±5.96 MPa, and 24.25±7.79 MPa, respectively) to that of the healthy cartilage; however, 9.1 w/v% IPN augments to that of healthy cartilage (38.98±8.61 MPa, **Fig. 3C**).

**Figure 3.**
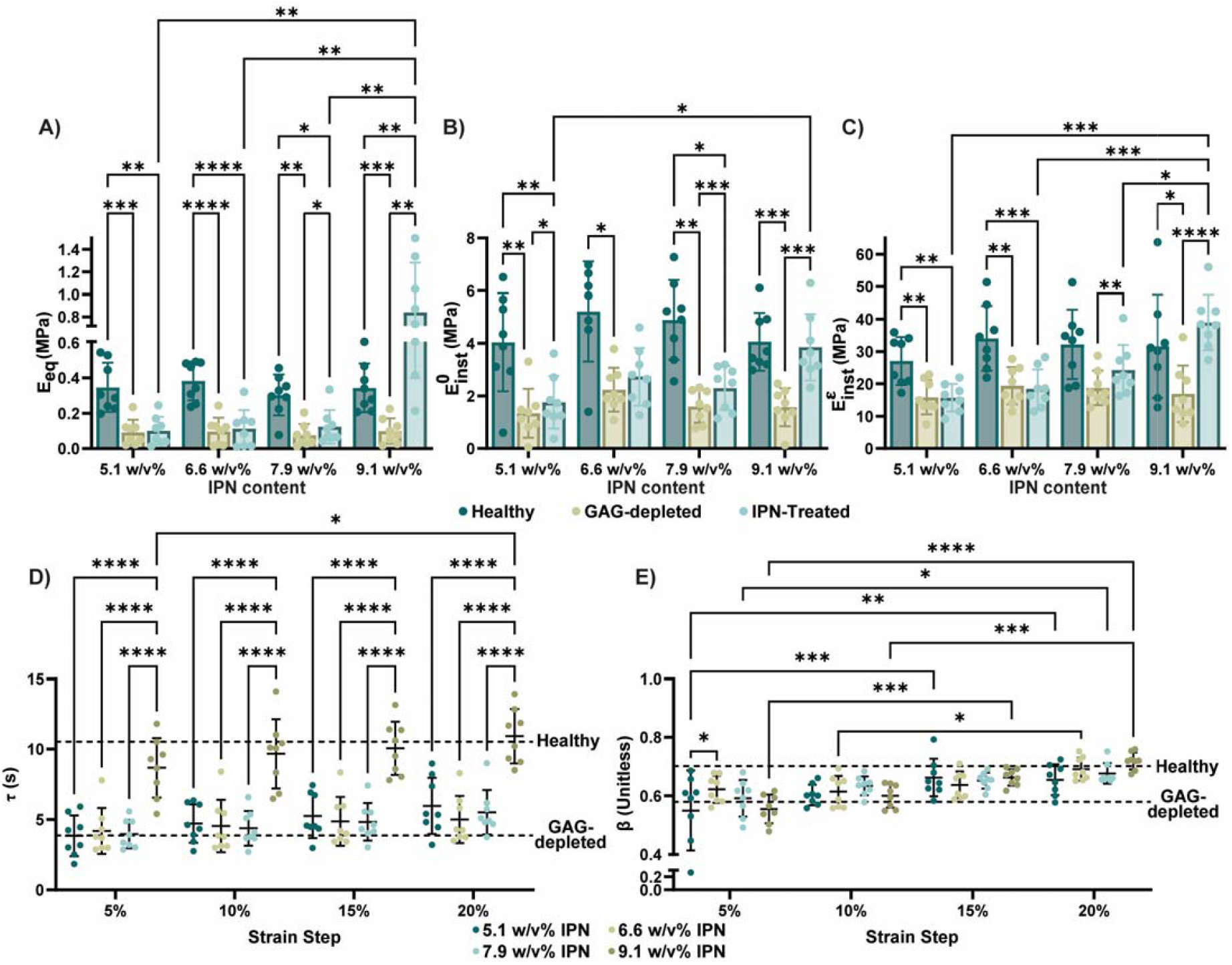
The mechanical properties of GAG-depleted cartilage are restored to healthy levels with the SPM IPN. The **A)** equilibrium modulus (), **B)** initial instantaneous modulus (), and **C)** strain-dependent instantaneous modulus () were determined for each cartilage sample as it progressed through the intact, GAG-depleted, and IPN-treated states identified by symbol color. The samples are grouped along the x-axis based on the concentration of IPN installed in the matrix. All three of these stiffness measurements decrease with GAG-depletion, and only the 9.1 w/v% restores them to healthy levels. n=7 or 8 cartilage samples per group. Data is represented as mean±S.D. Significance determined by 2-way paired ANOVA or mixed-effects model, ^*^ p<0.05, ^**^ p<0.01, ^***^ p<0.001, ^****^ p<0.0001. Similarly, the **D)** relaxation time (τ) and **E)** stretching parameter (β) of cartilage samples treated with the four IPN contents were calculated at each strain step of the stress relaxation procedure. The horizontal dashed lines represent the average values for intact and GAG-depleted samples. τ only returns to healthy levels with the 9.1 w/v% IPN, while β depends on strain level to a greater degree than IPN content and returns to that of intact cartilage at 20% strain. n=8 cartilage plugs per group. Data is represented as mean±S.D. Significance determined by 2-way unpaired ANOVA, ^*^ p<0.05, ^**^ p<0.01, ^***^ p<0.001, ^****^ p<0.0001.

The viscoelastic relaxation time (τ) and stretching parameter (β) significantly decrease by GAG depletion (**Fig. S3A-D and S4A-D**). After photopolymerization, IPN tissue concentrations of 5.1 w/v%, 6.6 w/v% and 7.9 w/v% do not restore τ to that of the intact cartilage across strain steps (at 20% strain: 5.97±2.0s, 5.005±1.67s, and 5.52±1.58s, respectively); however, 9.1 w/v% IPN restores τ to that of the healthy cartilage (at 20% strain: 10.93±2.95s), which is also significantly higher than the other IPN concentrations (**Fig. S3A-D and 3D**). β of IPN-treated cartilage increases with escalating strain, approaching that of the intact cartilage at 20% strain for all four IPN tissue concentrations: 0.66±0.052, 0.69±0.039, 0.68±0.035, and 0.72±0.029, respectively. β does not differ significantly between IPN tissue concentrations at each given strain level (**Fig. S4A-D and 3E**).

### 3.3 IPN restores the fixed negative charge density of GAG-depleted cartilage, integral to Donnan osmotic swelling pressure

By measuring the change in cartilage material properties tested in a low ionic strength solution compared to a high ionic strength solution, we used Donnan osmotic swelling to evaluate whether the 9.1 w/v% IPN restored the fixed negative charge density of the GAG-depleted cartilage to that of healthy cartilage. We calculated the ratios of 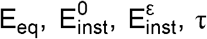, and β for cartilage plugs tested in 0 mOsm water and 400 mOsm saline for both healthy and IPN-treated cartilage. E_eq_characterizes the non-fibrillar matrix component contributing to cartilage stiffness (i.e. IFLS); therefore owing to osmotic swelling pressurization, for the same applied compressive load, the cartilage deforms 50% less in 0 mOsm water compared to 400 mOsm saline, indicating increased tissue stiffness, where the ratio 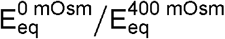 is 2.38±0.42 and 2.01±0.38, respectively, for healthy and IPN-treated cartilage (**Fig. 4A and S4A-B**). Since 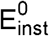 describes the modulus of the fibrillar component contributing to tissue stiffness at the onset of the strain application, and 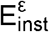 describes the magnitude of the change in the modulus of the fibrillar component contributing to tissue stiffness as the strain changes, these metrics are less affected by changes in ionic strength. The ratio 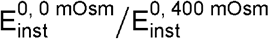 is 1.08±0.16 and 1.33±0.25 for healthy and IPN-treated cartilage, respectively (**Fig. 4B and S4C-D**). 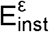 of the IPN-treated cartilage slightly decreases in low ionic strength conditions, where the ratio 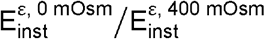 is 1.01±0.23 and 0.87±0.38, respectively, for healthy and IPN-treated cartilage (**Fig. 4C and S4E-F**), although, these values are not significantly different from each other.

**Figure 4.**
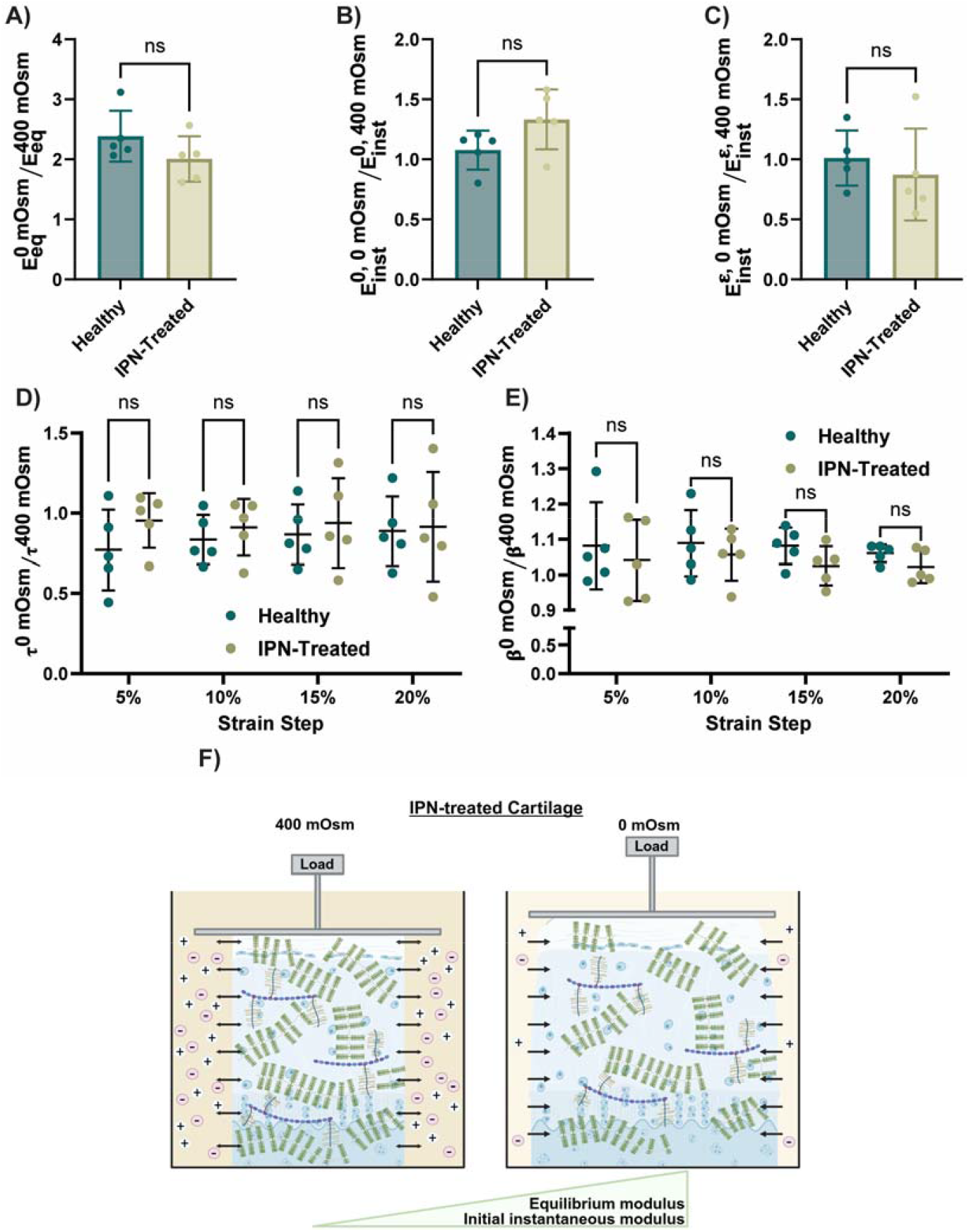
IPN restores fixed negative charge density to that of native GAG-network as measured by Donnan osmotic swelling pressure. To observe Donnan osmotic swelling pressure, healthy and IPN-treated cartilage samples underwent mechanical testing in 400 mOsm saline and 0 mOsm water. The ratios **A)**, **B)**, **C)**, **D)**, and **E)** were calculated to identify changes to the material properties between the solutions. **A-C)**, and are both greater than 1, showing the cartilage is stiffer in 0 mOsm water than 400 mOsm saline while remains close to 1. n=5 cartilage samples per group. Data represented as mean±S.D. Significance determined by unpaired t-test, NS=non-significant. **D and E)**, is less than 1 while is greater than 1 for healthy and IPN-treated cartilage, but there is no significant difference between intact and IPN-treated cartilage with these parameters. n=5 cartilage samples per group. Data represented as mean±S.D. Significance determined by 2-way unpaired ANOVA, NS=non-significant. **F)** A schematic representation of Donnan osmotic swelling pressure. Direction of water flow indicated by arrows for 400 mOsm and 0 mOsm. For the same applied compressive load, the cartilage deforms less in 0 mOsm water compared to 400 mOsm saline because of osmotic swelling pressurization, indicating increased tissue stiffness.

When tested in 0 mOsm saline compared to 400 mOsm saline, both the healthy and IPN-treated cartilage exhibit slightly lower relaxation times across strain levels, where τ ^0 mOsm^/ τ ^400 mOsm^ There is no difference between the healthy and IPN-treated cartilage **(Fig. 4D and S4G-H**). The stretching parameter, β, increases slightly when tested in 0 mOsm water compared to 400 mOsm saline for both the healthy and IPN-treated cartilage, where β ^0 mOsm^ / β ^400 mOsm^ > 1 across the four strain steps (**Fig 4E and S4I-J**).

The tissue stiffnesses of healthy and IPN-treated cartilage tested in neutral charge 400 mM maltose solution are nearly identical to those of cartilage plugs tested in 0 mOsm water, identifying that mobile ion transport affects the material properties of cartilage, and manifests as cartilage swelling in low ionic strength solutions (**Fig. S5**). These results confirm that the SPM-IPN reconstitutes the fixed negative charge density lost with GAG depletion (**Fig. 4F**).

Additionally, to assess the non-electrostatic contribution to cartilage mechanical properties provided by the GAG-network and IPN, we conducted mechanical testing in high tonicity saline (2000 mOsm) and compared the results to those acquired in 400 mOsm saline by taking the ratio of each parameter in 2000 mOsm aline to 400 mOsm saline. of both healthy and IPN-treated cartilage decrease in 2000 mOsm saline compared to 400 mOsm saline, as for healthy, and IPN-treated cartilage are 0.45±0.047 and 0.83±0.17, respectively (**Fig. S6A**). Similarly, is 0.70±0.12 and 0.82±0.19 for healthy and IPN-treated cartilage, respectively, demonstrating that decreases in high ionic strength solution when compared to 400 mOsm saline (**Fig. S6B**). 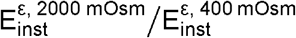 and 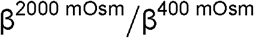 approximately 1 for both healthy and IPN-treated cartilage, while 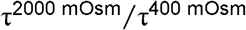 is ∼1 for healthy cartilage and ∼1. for IPN-treated cartilage across the four strain steps (**Fig. S6C-E**).

### 3.4 IPN reduces infiltration of inflammatory cytokines into GAG-depleted cartilage

Proinflammatory cytokines are elevated in OA synovial fluid. GAG-depletion renders cartilage susceptible to tissue partitioning of deleterious cytokines, enabling access to chondrocytes. Employing FITC-dextran as a visualizable surrogate for these inflammatory cytokines, IPN treatment reduces the permeability of FITC-dextran into the GAG-depleted tissue matrix. After incubating cartilage plugs for 24-hours in FITC-dextran solution, fluorescent imaging demonstrates elevated fluorescent signal in GAG-depleted cartilage compared to healthy cartilage; however, IPN-treated cartilage exhibited low fluorescent levels, similar to the healthy cartilage (**Fig. 5A**). Partition coefficients of FITC-dextran into cartilage increase 7-fold from 0.024±0.0084 for healthy cartilage to 0.17±0.044 for GAG-depleted cartilage (**Fig. 5B**). IPN treatment significantly decreases the partition coefficient by more than 50% to 0.079±0.053, albeit not equivalent to that of healthy cartilage (**Fig. 5B**).

**Figure 5.**
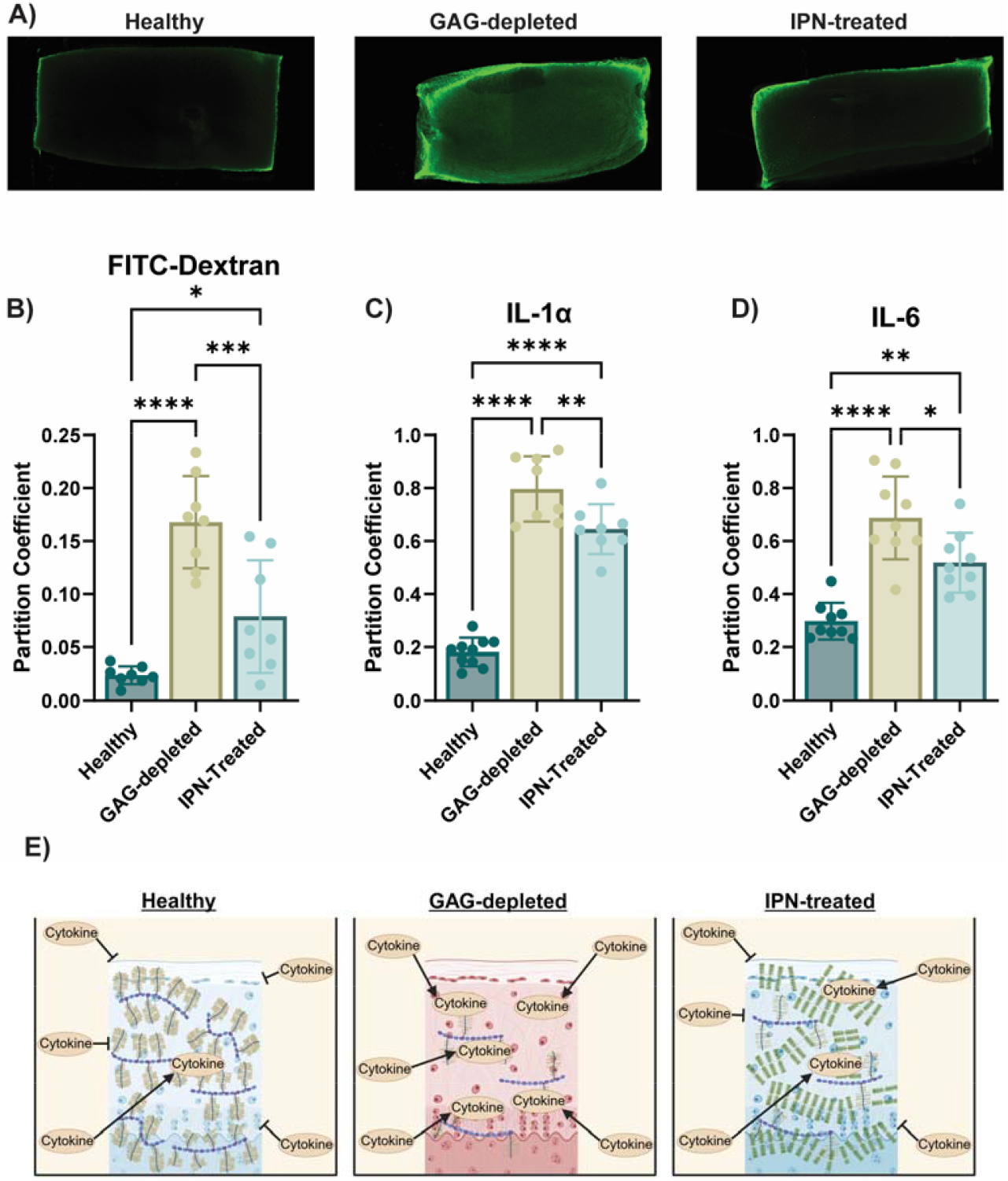
IPN treatment reduces cytokine infiltration into the cartilage matrix, restoring the excluded volume effect to GAG-depleted cartilage. **A)** Representative confocal images of samples incubated in solutions of FITC-dextran show more solute enters GAG-depleted cartilage compared to intact and IPN-treated cartilage. **B)** The FITC-dextran partition coefficient reveals significantly more infiltration of FITC dextran into GAG-depleted cartilage than into intact or IPN treated GAG-depleted cartilage. n=8 cartilage samples per group. Data is represented as mean±S.D. Significance determined by 1-way unpaired ANOVA, ^*^ p<0.05, ^***^ p<0.001, ^****^ p<0.0001. The partition coefficients of **C)** IL-1α and **D)** IL-6 in GAG-depleted cartilage are significantly higher compared to intact and IPN-treated cartilage. n=8, 9 or 10 cartilage samples per group. Data is represented as mean±S.D. Significance 1-way unpaired ANOVA: ^*^ p<0.05, ^**^ p<0.01, ^***^ p<0.001, ^****^ p<0.0001. **E)** Graphical representation of IPN treatment restoring the excluded volume effect of GAG-depleted cartilage, thereby preventing cytokine infiltration into the matrix similar to intact cartilage.

We next measured the partition coefficient of IL-1α, an anionic cytokine integral to the onset and progression of OA.[69] The partition coefficient increases more than 4-fold to 0.80±0.12 after GAG depletion compared to 0.18±0.053 for healthy cartilage; IPN treatment lowers the partition coefficient for GAG-depleted cartilage by 23% to 0.65±0.094 (**Fig. 5C**). Similarly, the partition coefficient of IL-6, a potent inflammatory cytokine upregulated in OA, [70] increases more than 2-fold to 0.69±0.16 after GAG-depletion, compared to 0.3±0.070 for healthy cartilage; IPN treatment reduces the partition coefficient of GAG-depleted cartilage by 25% to 0.52±0.11 (**Fig. 5D**). Therefore, IPN treatment reduces the permeability of interleukins into GAG-depleted cartilage (**Fig. 5E**).

### 3.5 IPN reconstitutes degenerated cartilage tissue modulus

Using osteochondral plugs excised from the tibial plateau of mature bovine stifle joints (**Fig. S7A**), we non-destructively assessed the biochemical composition of the tissue (i.e., regression coefficients GAG_score,_ Collagen_score_, Water_score_, Bone_score_ and SPM_score_ portray the relative contribution of each constituent to the composite Raman tissue spectra) by Raman spectroscopy. Therefore, the SPM_score_ is used as a surrogate measurement for the amount of the SPM-IPN within the cartilage matrix. We also measured tissue stiffness by indentation, before and after installation of the SPM-IPN into the degenerated cartilage. Raman spectroscopy shows that the GAG levels were lowest in the articular tissue adjacent to the visibly degenerated (Outerbridge 2) cartilage lesion (**Fig. S7B**).[66, 67] Following installation of the SPM-IPN into the degenerated cartilage, SPM is detected in all articular tissue as seen with the large increase in SPM_score_ in the Post-IPN group compared to the Pre-IPN group (**Fig. 6A**). The average SPM_score_ across all of the osteochondral samples increases 20-fold to 0.08±0.014 after IPN treatment compared to 0.0041±0.0017 prior to IPN treatment (**Fig. 6B**). The SPM-IPN stiffens the degenerated tissue by nearly 50% from 0.45±0.096 MPa to 0.66±0.13 MPa (**Fig. 6 C-D**). With IPN treatment, the average Water_score_ significantly decreases, while the Collagen_score_ remains unchanged (**Fig. S7C and D**). These data suggest that the SPM-IPN reduces tissue porosity (as represented by water content) by insinuating into the tissue pores thereby decreasing pore volumes while relatively increasing the local fixed negative charge density that together retard interstitial fluid transport to reconstitute the IFLS of the degenerated cartilage.

**Figure 6.**
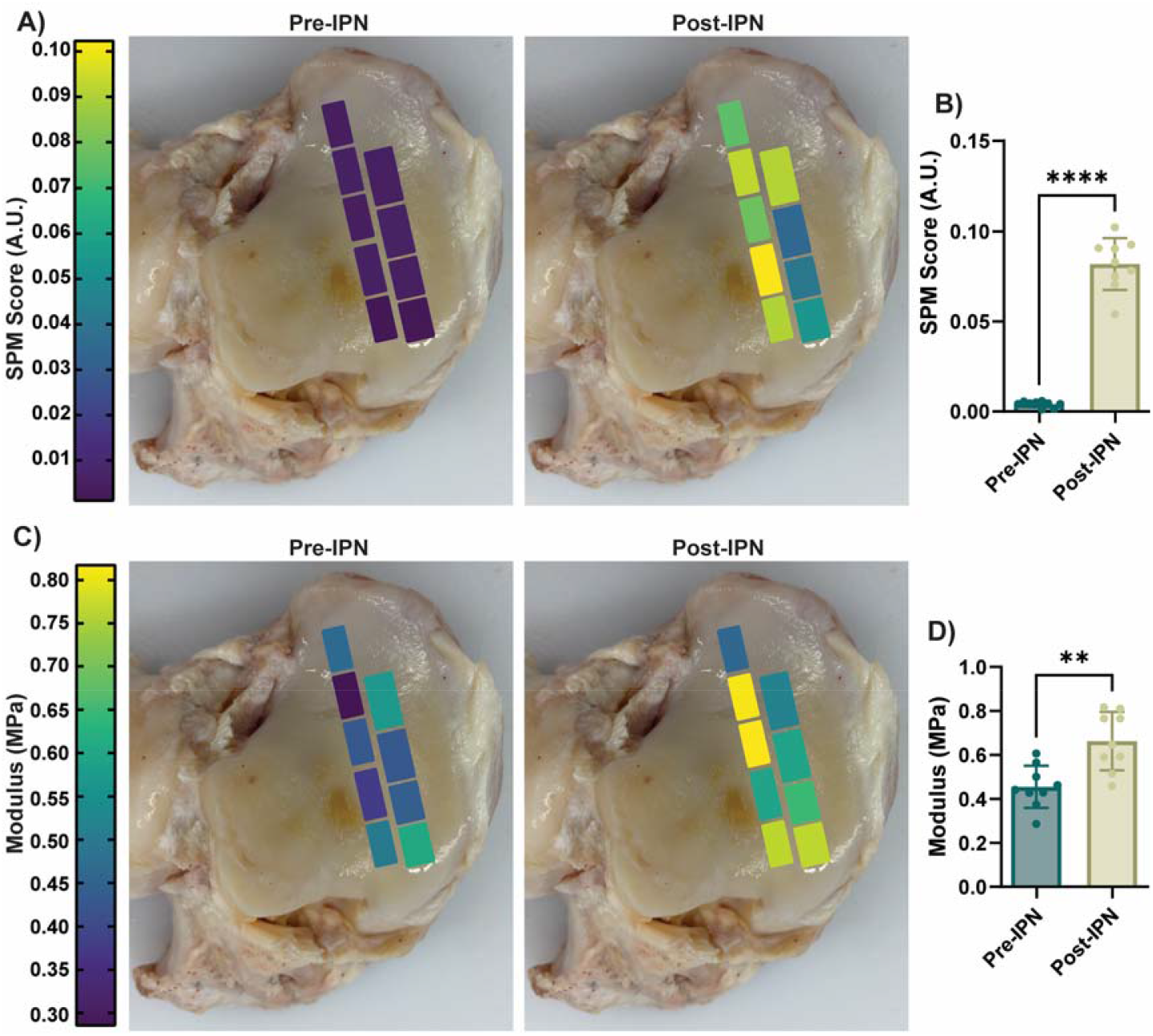
SPM-IPN treated arthritic bovine cartilage is detectable in the tissue by Raman spectroscopy and increases tissue stiffness. Heat map of **A)** SPM score determined by Raman spectroscopy and **C)** tissue modulus for individual osteochondral plugs before and after IPN treatment overlaid onto the tibial plateau corresponding to where samples were harvested. **B)** Average SPM score is significantly greater in the post-IPN treatment group compared to the pre-IPN group. n=9 samples per group. Data is represented as mean±S.D. Significance determined by paired t-test. ^****^ p<0.0001. **D)** Average modulus is significantly stiffer after IPN treatment; n=9 samples per group. Data represented as mean±S.D. Significance determined by paired t-test. ^**^ p<0.01.

## 4. Discussion

ECM composition governs the material properties of hyaline cartilage requisite for articular joint function: support high compressive loads with low sliding friction, while retarding infiltration of inflammatory cytokines that provoke cartilage degradation. The porous-permeable non-fibrillar matrix coordinates water via highly sulfated, anionic, GAGs decorating a core protein that complexes with hyaluronic acid, forming a bottlebrush structure, complemented by a COLII fibrillar network that provides structural support and tensile strength to resist Donnan osmotic swelling pressure. The progressive loss of major constituents comprising the cartilage ECM is characteristic of joint disease. GAG depletion from the ECM reduces *IFLS* and transfers load to the COLII fibril network, which subsequently breaks down, culminating in increased hydraulic permeability, and decreased cartilage stiffness. Decreased GAG content and increased tissue porosity reduce the excluded volume effect. Diminished repulsion of anionic solutes through charge interactions allows proinflammatory cytokines in the synovial fluid to permeate the GAG-depleted ECM and access chondrocytes.

While considerable research focuses on tissue engineering strategies that physically replace damaged cartilage with a polymer scaffold loaded with chondrogenic cells that produce a neocartilage, strategies to replace or augment lost GAGs with synthetic alternatives that reconstitute the tissue’s ECM are less studied. Typically, GAG mimetic strategies rely on the functionalization of naturally derived carbohydrates that lack effective mechanisms for long-term retention within the cartilage ECM.[71-75] Our synthetic approach recapitulates the naturally occurring proteoglycan network via an IPN produced using SPM, PEGDa, and a photoinitiation system consisting of TEOA, NVP, and eosin Y. Analogous to GAGs, SPM imparts anionic charge via a sulfonate group. Crosslinks provided by PEGDa ensure that the IPN interlinks with the existing collagen matrix. The photoinitiation system provides control over the initiation of the free radical polymerization process, since light must be introduced to the system for the polymerization to begin, to avoid premature network formation, which is a risk with chemical initiators that spontaneously form radicals.[76]

The concentration of the SPM monomer in the treatment bath dictates the amount of SPM polymer formed in the cartilage. The 5 w/v%, 20 w/v%, 40 w/v% and 60 w/v% SPM monomer bath concentration result in 5.1, 6.6, 7.9, and 9.1 w/v%. Of the concentrations, only the 9.1 w/v% concentration significantly increases the stiffness, relaxation time, and stretching parameter of the GAG-depleted tissue to that of healthy cartilage, indicating that there is a minimum amount of anionic polymer required to reconstitute the material properties of degenerated cartilage. The 9.1 w/v% IPN restores both E_eq_, reflecting the porous-permeable hydrogel component of cartilage, and 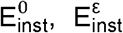 reflecting the fibrillar component of the cartilage ECM, thereby demonstrating entanglement of the polymer with the native COLII network via light activated polymerization of the SPM and PEGDa.

The fixed negative charge density of ECM cartilage is critical to cartilage function. Van Mow’s triphasic theory describes articular cartilage swelling (and IFLS) dependency on the magnitude and distribution of the fixed negative charge density provided by GAGs within the ECM, the stiffness of the biphasic collagen-proteoglycan matrix, and the ionic concentration of mobile cations and anions in the bathing solution.[77] Since the negative charge groups are impermeable, and cannot change, only interstitial water and the mobile counter-ions (Na^+^ cations) of the saline bath change. The equilibrium equation for the mobile ions in the saline bathing solution and for the interstitial water are expressed in terms of their chemical potentials whose gradients are the driving forces for their movements in and out of the cartilage ECM. These chemical potentials depend on osmotic fluid pressure, salt concentration, solid matrix dilatation, and fixed charge density. Accordingly, cartilage submerged within a bathing solution reaches a (Donnan) equilibrium state of electrochemical charge separation (Donnan potential), osmolarity, and mechanical stress (for compressed cartilage). In a solution of low ionic strength, the fixed negative charge groups in the cartilage ECM drives fluid flow (mobile cations and water) into the matrix which pressurize and stiffen the tissue (Donnan osmotic swelling pressure).[78, 79] Thus, we used Donnan osmotic swelling to confirm that SPM-IPN reconstitutes the compressive stiffness of GAG-depleted cartilage by comparing the material properties of cartilage tested in water (0 mOsm) and in 400 mOsm saline (physiologic ionic concentration and osmolarity for cartilage and synovial fluid). E_eq_portrays the porous-permeable hydrogel component of cartilage ECM most affected by different ionic strength bathing solutions, while 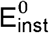, and 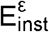 reflect the collagen-fibrillar component of the cartilage ECM that is relatively unaffected by different ionic strength solutions. In water, E_eq_for both healthy and IPN-treated cartilage doubles relative to the E_eq_in 400 mOsm saline, validating that SPM-IPN augments the fixed negative charge density of the GAG-depleted ECM. Further, to determine if the SPM negatively charged groups are responsible for tissue swelling and not the influx of mobile ions into the tissue to achieve Donnan ion equilibrium, we measured the stiffness of the cartilage plugs in an uncharged iso-osmotic solution. The stiffness values of cartilage plugs submerged in 400 mM maltose is comparable to the stiffnesses values of cartilage plugs measured in 400 mOsm saline and water. E_eq_are equivalent for both the healthy and IPN-treated cartilage tested in 400 mOsm maltose and water, establishing that the fixed negative charge groups of SPM, similar to GAG, are responsible for tissue swelling. Finally, the mechanical properties of healthy and IPN-treated cartilage were determined with the tissue submerged in a high ionic strength solution (2000 mOSm), which leads to a shielding of the charged matrix as the mobile cations neutralize the fixed negative charge density provided by the GAG network and IPN, respectively. Therefore, cartilage in the high ionic strength solution represents the non-electrostatic contribution of the GAG network and the IPN to the cartilage mechanical properties.[80] Taking the ratio of in 2000 mOsm saline to 400 mOsm saline indicates the non-electrostatic contribution of the GAG network and IPN to the equilibrium stiffness is 45% and 83%, respectively. The increased non-electrostatic contribution of the IPN to suggests that the solid content of the IPN dictates a majority of the equilibrium properties of the IPN-treated cartilage, while the non-electrostatic contribution is comparable stiffness to the electrostatic contribution of the GAG network to the equilibrium stiffness in healthy cartilage. The non-electrostatic contribution of the GAG network contributes to 70% and 82% of in healthy and IPN-treated cartilage, respectively. describes the tension the fibrillar components of the cartilage matrix are under at the onset of strain application, therefore, it is not unsurprising that the non-electrostatic contribution of the GAG network is higher for this mechanical property com ared to of healthy cartilage. The non-electrostatic contribution of the IPN largely determines, similarly to, demonstrating the solid content of the IPN has a significant impact on the stiffness properties of the treated cartilage.

Both the intact and IPN-treated cartilage exhibit slightly lower relaxation times across strain levels when tested in water compared to 400 mOsm saline. Similarly, the IPN-treated cartilage tested in 2000 mOsm saline demonstrates slightly higher relaxation times across strain levels compared to in 400 mOsm saline. Tissue swelling in low ionic strength solutions may increase pore size and permeability allowing unbound interstitial water to more quickly exit the matrix during compressive loading thereby shortening the relaxation time. Our observation of shorter τ in low ionic strength solutions differs from a previous report that in low ionic strength solution cartilage had a slightly longer τ when compared to a high ionic strength solution; however, this report supports our findings that β is larger when tests are conducted in a low ionic strength solution.[64]

The inflammatory environment of an osteoarthritic joint leads to increased cytokines in the synovial fluid. Increased tissue porosity and decreased GAG content reduce the excluded volume effect through charge interactions, diminishing repulsion of anionic solutes [35] and allowing cytokines in the synovial fluid to permeate the GAG-depleted ECM to gain access to chondrocytes. The cytokines promote a phenotypic transition of chondrocytes to produce catabolic enzymes such as matrix metalloproteinases that exacerbate cartilage degradation. FITC-dextran, IL-1α, and IL-6 readily infiltrate the GAG-depleted cartilage. Albeit not equivalent to that of healthy cartilage, IPN treatment of the GAG-depleted cartilage partly retards partitioning of these solutes into the ECM in a size dependent manner, where IL-1α with a molecular weight of ∼18 kDa, exhibits a larger partition coefficient than IL-6 with a molecular weight of ∼21 kDa. This result differs from healthy cartilage where the partition coefficient for IL-6 is larger than that for IL-1α. Following partitioning of FITC-dextran into the IPN-treated cartilage, the fluorescent images show high concentrations of dextran at the periphery of the cartilage sample, with very low concentrations in the bulk of the tissue. Thus, while IPN treatment of GAG-depleted cartilage might reduce partitioning of cytokines into the core of the tissue, accumulation at the surface renders superficial zone cartilage at particular risk.

To establish clinical relevance, in addition to using enzymatically GAG-depleted bovine cartilage, we examined the mechanism of action and effectiveness of the SPM-IPN in naturally occurring degenerated cartilage using osteochondral plugs excised from visibly degenerated (Outerbridge 2) mature bovine tibial plateau. We conducted non-destructive mechanical and compositional assessments before and after IPN installation. SPM-IPN treatment stiffens the degenerated tissue by nearly 50%. We used our Raman spectroscopy-based platform to non-destructively appraise the composition of degenerated cartilage augmented with SPM IPN.[66, 67] We decomposed the Raman spectra of the SPM-IPN infused cartilage composite into the “fingerprint” spectra of individual ECM constituents -GAGs, collagen, water, and SPM, where the derived multivariate regression coefficients that optimize the “fit” of the constituent spectra to the overall “fingerprint” profile represents the proportional contribution of each constituent to the cartilage composite. After SPM infusion and light-activated polymerization, SPM-IPN is present in all articular tissue; the average SPM_score_ increases 20-fold, the average Water_score_ significantly decreases, while the Collagen_score_ remaines unchanged. These data suggest mechanistically that SPM-IPN insinuates into the pores, decreasing tissue porosity (since water is located within the pore space of the matrix, water content is directly related to tissue porosity). The reduction in pore size combined with the relative increase in local fixed negative charge density act together to retard interstitial fluid flow, thereby reconstituting IFLS to degenerated cartilage.

While the data strongly support the effectiveness of SPM-IPN to reestablish *IFLS*, there are limitations in the current study. We used both enzymatically degraded immature bovine cartilage plugs and naturally degenerated mature bovine cartilage to simulate osteoarthritis pathoanatomy. While bovine cartilage is an efficient model to establish feasibility, studies using human tissues of various osteoarthritic states are needed to validate our approach. Studies are required to evaluate chondrocyte viability *in vivo in a* post-traumatic OA large animal model to examine how IPNs affect the entire diarthrodial synovial joint, with attention to off-target tissue effects. Operationally reconstituting GAG-depleted cartilage using a synthetic sulfonated interpenetrating polymer to reestablish *IFLS* will require a minimally invasive, intra-articular procedure during conventional arthroscopy: after correcting the internal derangement contributing mechanically to acute or chronic chondrocyte injury and tissue wear, the joint is irrigated with saline to neutralize catabolic enzymes that degrade cartilage; SPM is then instilled into the joint and polymerized with white light to reconstitute damaged cartilage material properties. The white light source is commonly used in arthroscopes, and the eosin Y based system has been used successfully in the clinic with a regulatory approved product for sealing lung tissue lesions.[59]

## 5. Conclusions

We restore the material properties of damaged cartilage critical to diarthrodial joint function by forming an IPN with the native collagen using a synthetic, hydrophilic, and biocompatible GAG-mimetic polymer to restore *IFLS* and reestablish the integrity of the collagen fibril network. For the IPN, the monomer 3-sulfopropylmethacrylate (SPM) and crosslinker polyethylene glycol diacrylate (PEGDA) form a sulfonated and anionic polymer network that entangles with the existing collagen matrix upon visible photolysis. Practically, this approach is adaptable to minimally invasive arthroscopy as the pre-SPM IPN material can be delivered to the joint and subsequently polymerized using the white light scope. Using both enzymatically GAG-depleted and naturally degenerated cartilage as models of early OA, the SPM-IPN restores the stiffness of degraded cartilage by filling the porous tissue with a dense highly sulfonated, anionic hydrogel that retards water transport to restore tissue *IFLS*. Additionally, the SPM-IPN reduces the infiltration of inflammatory cytokines that upregulate catabolic matrix metalloproteinases and downregulate GAG production. Currently, there are no treatments that restore the material properties of hyaline cartilage tissue critical to its mechanical function and impart chondroprotection after OA induction. This work provides a strategy for treating early OA by reestablishing *IFLS* in GAG-depleted cartilage using a synthetic sulfonated interpenetrating polymer and a framework to design new in-situ polymerizing biomaterials to repair damaged ECM in tissues.

## CREDIT authorship contribution statement

**Christian D. DeMoya:** Conceptualization, Methodology, Formal analysis, Investigation, Data Curation, Writing – Original Draft, Writing – Review & Editing, Visualization, Project administration. **Pierson Husted:** Investigation, Formal analysis. **Dev R. Mehrota:** Methodology, Formal analysis, Investigation. **Michael B. Albro:** Resources, Writing – Review & Editing, Supervision. **Brian D. Snyder:** Conceptualization, Resources, Writing – Review & Editing, Supervision. **Mark W. Grinstaff:** Conceptualization, Resources, Writing – Original Draft, Writing – Review & Editing, Supervision, Funding acquisition.

## Acknowledgements

The authors would like to acknowledge Matthew Taylor for his assistance with capturing the fluorescent images presented in this work. The content is solely the responsibility of the authors and does not necessarily represent the official views of the National Institute of Health.

## Declaration of Generative AI and AI-assisted Technologies in the Writing Process

During the preparation of this work the author(s) used no tools or services.

## TOC Figure

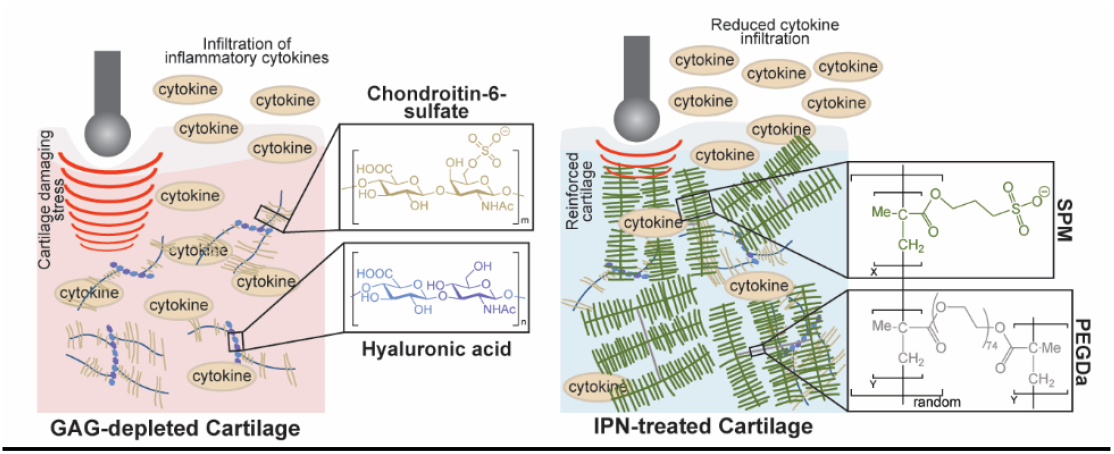

